# Abnormal Tau Protein Influences Intercellular Mitochondrial Transfer Between Astrocytic and Neuronal Cells

**DOI:** 10.1101/2025.04.11.648113

**Authors:** Aurélien Riou, Aline Broeglin, Andreas Papassotiropoulos, Anne Eckert, Amandine Grimm

## Abstract

Tauopathies are a group of neurodegenerative diseases characterized by the pathological accumulation of abnormal tau protein. A consequence of tau pathologies is mitochondrial dysfunctions, which affect essential processes such as mitochondrial transport, bioenergetics, and dynamics. Given the high energy demands of neurons, tau-induced mitochondrial impairment significantly contributes to neuronal vulnerability and degeneration. Recent studies have revealed that cells can transfer mitochondria between them to help energy-deficient cells. This process, known as intercellular mitochondrial transfer, occurs through two different pathways: an indirect transfer via extracellular vesicles and a direct transfer via tunneling nanotubes and gap junctions. Given the known impact of abnormal tau protein on mitochondrial transport and actin filament dynamics, we hypothesized that intercellular mitochondrial transfer could be altered in the context of tauopathies. Therefore, this study aimed to investigate mitochondrial transfer between astrocytic and neuronal cells and assess how abnormal tau protein may influence this process. Our results showed that abnormal tau protein enhances mitochondrial transfer from astrocytic to neuronal cells. Notably, this transfer occurs mainly via contact-dependent mechanisms. In both pathological and healthy conditions, the transferred astrocytic mitochondria either fused with the mitochondrial network of recipient cells or were degraded in the lysosomes or remained isolated in the cytosol. Our data highlight a novel pathway by which abnormal tau protein impacts mitochondrial function, namely the transfer of astrocytic mitochondria to neuronal cells.

## Introduction

The brain is a high-precision electro-chemical organ that consumes approximately 20% of the body’s energy, although comprising only 2% of total body mass, to sustain its functions [1]. To meet this high energy demand, the brain relies on mitochondria, essential organelles in all eukaryotic cells, because of their critical role in adenosine triphosphate (ATP) production via oxidative phosphorylation, the primary energy-generating pathway in cells [2]. Multiple studies have demonstrated that brain energy production is impaired in neurodegenerative diseases, including tauopathies, thereby contributing to disease progression [3,4].

Tauopathies encompass a heterogeneous group of neurodegenerative disorders, such as Alzheimer’s disease (AD) or frontotemporal lobar degeneration (FTLD), characterized by the accumulation of abnormal tau protein [5]. Clinically, these diseases manifest with a broad spectrum of neurological deficits, including cognitive and psychiatric disorders, motor dysfunctions, and difficulties with speech, swallowing, and vision [6]. Unfortunately, no cure has been identified for these conditions yet. Physiologically, tau is a member of the microtubule-associated protein (MAP) family [1]. It is predominantly expressed in neurons, especially within axons, where it plays a pivotal role in stabilizing microtubules by binding to tubulin and promoting its polymerization. Consequently, tau is essential for the axonal transport of organelles, neuronal plasticity, axon growth, elongation, morphogenesis, differentiation, and neurite polarity [1,7,8]. In tauopathies, tau becomes abnormally hyperphosphorylated, leading to its aggregation, causing microtubule destabilization and, ultimately, neuronal dysfunction and death [9]. Besides, a prominent hallmark of all these tau-related diseases is the impairment of brain bioenergetics and mitochondrial function [10,11]. Indeed, abnormal tau protein is known to impair almost all mitochondrial functions, namely mitochondrial transport [12,13], dynamics [14], bioenergetics [3,4] and mitophagy [15], thereby contributing to neuronal degeneration. However, the precise molecular mechanisms by which tau affects mitochondrial function remain elusive.

Recently, studies on intercellular mitochondrial transfer (IMT) have emerged, demonstrating this mechanism across various organs and pathological contexts [16]. Studies have shown that healthy cells can transfer mitochondria to support dysfunctional cells, while damaged cells may pass defective mitochondria to healthy cells for degradation and recycling [2]. IMT has been observed in various cell types, including astrocytes, mesenchymal stem cells, and immune cells, highlighting its broad physiological and pathological significance. Two primary mechanisms for intercellular mitochondrial transfer have been identified:

- Extracellular vesicles (EVs): EVs are bilayer membranous structures secreted into the extracellular space. Astrocytes have been shown to shed EVs containing functional mitochondria [17].
- Tunneling Nanotubes (TNTs): TNTs are membrane channels composed of F-actin that connect two or more cells. This channel allows the transfer of cytoplasmic molecules or organelles from one cell to another [18].

Furthermore, other mechanisms may be possible as for example extracellular mitochondria internalization by recipient cells or transferred via gap junctions [19].

Astrocytes are well known for their crucial supportive role in the central nervous system, particularly for neurons. They are known to regulate the neuronal microenvironment by providing metabolic substrates, clearing oxidative stress, and modulating synaptic activity [20-22]. Given their essential support functions and substantial mitochondrial pool, despite their primarily glycolytic metabolism [23], astrocytes have been studied for their involvement in IMT and identified as key players in mitochondrial transfer to neurons [17,24-26].

Evidence has been provided that intercellular mitochondrial transfer (IMT) between astrocytes and neurons has beneficial effects on the brain, such as increasing ATP levels and neuronal viability in recipient cells in brain injury [17,25,26].

Mitochondrial transport and functions are known to be negatively impacted in the context of tauopathies [1,27] However, no study has explored whether abnormal tau protein impacts IMT between neurons and astrocytes. Determining whether astrocytes can transfer healthy mitochondria to neurons affected by tau pathology could provide critical insights into neuroprotection and potential therapeutic strategies. Therefore, this study investigated whether mitochondrial transfer occurs between astrocytes and neurons in the presence of disease-associated tau protein. We also aimed to identify which transfer pathway was involved and to determine the fate of the transferred mitochondria in recipient cells.

### Culture, differentiation, and co-culture of neuronal and astrocytic cells

Human neuroblastoma SH-SY5Y cells, a widely used and well-established neuronal model, were purchased from ATCC® (CRL-2266™). The SH-SY5Y overexpressing the tau mutation P301L (P301L) tagged to a green fluorescent protein (GFP) were generated using lentiviral gene transfer [28,29] and kindly provided by the laboratory of Jürgen Götz (Queensland Brain Institute, University of Queensland, Brisbane, Australia). Human glioblastoma A172 cells, a well-established astrocytic model, were purchased from ATCC® (#CRL-1620™).

Both neuronal and astrocytic cell lines were cultured in Dulbecco’s Modified Eagle Medium (DMEM) (Sigma-Aldrich, #D6429) supplemented with 1% (v/v) penicillin/streptomycin (Bioconcept, #4-01F00-H), 1% GlutaMax (ThermoFisher Scientific, #35050061), 10% of heat-inactivated fetal bovine serum (FBS) (Corning, #35-015-CV). In addition, the SH-SY5Y cells culture medium was supplemented with 5% of heat-inactivated horse serum (Bioconcept, #2-05F00-I). To maintain stable expression of the P301L mutation and fluorescent tags, the culture medium was supplemented every two passages with 4.5 μg/mL of blasticidin (InvivoGen, #Ant-bl-1) for the GFP-tagged cells, while 500 μg/mL of geneticin sulfate (Santa Cruz Biotechnology, #SC-29065B) was used for cells stably expressing mitochondria-labeled RFP, or BFP2 tags. The cells were cultured in 10 cm^2^ dishes, split twice a week, and replated upon reaching approximately 80% confluence.

Prior to the experiments, SH-SY5Y-GFP cells bearing or not the P301L tau mutation were differentiated according to the following protocol. The day after plating, the culture medium was replaced with DMEM supplemented with 1% (v/v) penicillin/streptomycin, 1% GlutaMax, 1% nonessential amino acid (Gibco, #11140-035), 3% FBS, and 10 μM of retinoic acid (ThermoFisher Scientific, #04454002). After 48 hours, the medium was replaced with fresh DMEM supplemented with 1% (v/v) penicillin/streptomycin 1% GlutaMax, 1% FBS and 10μM of retinoic acid. Twenty-four hours later, the medium was changed to Neurobasal medium (ThermoFisher Scientific, #21103049) supplemented with 1% (v/v) penicillin/streptomycin, 1% of Glutamax, 1% of B27 (ThermoFisher Scientific, #17504044) and 50 ng/mL of BDNF (PeproTech, #450-02-100UG). The cells were then incubated in this medium for 48 hours before being used for co-culture. Forty-eight hours prior to the co-culture, the A172 cells were placed in Neurobasal medium supplemented with 1% (v/v) penicillin/streptomycin, 1% of Glutamax and 2% of B27.

### Lentivirus preparation and cell transduction

To study mitochondrial transfer from astrocytic to neuronal cells, we used transduced cells expressing GFP tags and fluorescent mitochondria. For this purpose, we employed custom-designed lentiviral vectors from VectorBuilder. These vectors can be identified using the following IDs, which provide detailed information on the VectorBuilder website (vectorbuilder.com).

1. **Green Fluorescent Protein (GFP)-labeled cells**: pLV[Exp]-Bsd-CMV>EmGFP (ID: VB231221-1477jqh).
2. **Mock vector**: pLV[Exp]-CMV>ORF_Stuffer (ID: VB900140-2612kkt).
3. **Red Fluorescent Protein-labeled mitochondria (mitoRFP)**: pLV[Exp]-CMV>{COX8 presequence:TagRFP-T}-{SV40_promoter}>Neo (ID: VB230714-1048hhg).
4. **Blue Fluorescent Protein 2 -labeled mitochondria (mitoBFP2)**: pLV[Exp]-CMV>{COX8 presequence:TagBFP2}-{SV40_promoter}>Neo (ID: VB240506-1616nnn).

SH-SY5Y and A172 cell lines were transduced at a multiplicity of infection (MOI) of 5, following the manufacturer’s protocol. Transduction efficiency was then confirmed by fluorescence imaging.

### Direct and indirect co-culture

Two co-culture methods were employed: direct and indirect co-culture using 3 μm mesh inserts (cellQART, #9313002) to simulate non-contact-dependent mitochondrial transfer. Both methods were conducted at a 1:1 ratio of SH-SY5Y to A172 cells for 24 or 48 hours.

### Fluorescent microscopy and colocalization analysis

After 24 or 48 hours of co-culture on precoated collagen coverslips (Neuvitro, #GC-18-1.5), cells designated for fluorescent microscopy were fixed with 4% paraformaldehyde (Sigma-Aldrich, #P6148-50G, CAS number: 30525-89-4) for 10 minutes at room temperature and washed three times with PBS. Slides were then mounted with an antifade mounting medium (Vectashield, #H-1000). Images were captured using a Nikon Eclipse Ti2 fluorescence microscope and deconvolved using Huygens software (version 24.10.0, Scientific Volume Imaging B.V.). To quantify the proportion of transferred mitochondria that are either degraded by recipient cells or integrated into their mitochondrial network, we performed a 48-hour direct co-culture of GFP-tagged SH-SY5Y (vector and P301L) expressing also the mitoRFP, with A172 expressing mitoBFP2. After the co-culture, lysosomes were stained with LysoTracker DeepRed (100 nM, 1 hour) (Invitrogen, #L12492) prior to cell fixation. To determine the fate of astrocytic mitochondria in neuronal cells, colocalization analysis was performed between A172 mitoBFP2 and either SH-SY5Y mitoRFP or LysoTracker-labeled lysosomes in SH-SY5Y vector/P301L cells. Colocalization was quantified using the JACoP plugin version in FIJI software (version 2.16.0), based on Manders’ coefficients. To generate 3D representations of each channel, false colors were used with the surface creation tool in Imaris (version 9.9.1, Oxford Instruments), namely SH-SY5Y cells are shown in grey, A172 mitoBFP2 in cyan, SH-SY5Y mitoRFP in magenta, and lysosomes in green. For each channel, surfaces were manually defined based on the visible signal in the z-stack images, following the structures observed by eye to outline the regions of interest.

### Flow cytometry

Flow cytometry experiments were performed to quantify IMT from A172 expressing mitoRFP to GFP-tagged SH-SY5Y (vector and P301L). We used an unstained (mock) control co-culture and fluorescence-minus-one (FMO) controls to determine the optimal voltages, compensation parameters, and gating strategy. After 24 or 48 hours of direct and indirect co-culture, cells designated for flow cytometry analysis were dissociated into single cells using Accutase (Chemie Brunschwig, #ICTAT104) and resuspended in HBSS (Gibco, #14065049). To eliminate potential cell aggregates, the suspension was passed through a 40 μm mesh filter (pluriSelect, #43-10040-60). Before analysis on the BD LSRFortessa flow cytometer, dead cells were stained with DAPI (Sigma, #MBD0015) at 1 μg/mL concentration. Data analysis was performed using Flowjo software (BD Biosciences, version 10.10.0) to assess the presence of A172 mitoRFP in GFP-positive SH-SY5Y cells.

### Statistical analysis

Statistical analyses were performed using GraphPad Prism software (GraphPad Software, version 10.4.1). All quantified data were presented as mean ± standard error of the mean (SEM). Data was submitted to a Grubbs test to determine and remove significant outliers. Statistical significance between two groups was assessed using Mann-Whitney test. A *p*-value of less than 0.05 was considered statistically significant, with significance levels denoted as follows: *p* < 0.05 (*), *p* < 0.01 (**), and *p* < 0.001 (***).

## Results

### Mitochondrial transfer occurs from astrocytes to neurons in both healthy and P301L-expressing cells

SH-SY5Y GFP-tagged cells, either carrying the P301L mutation or the empty vector, were co-cultured with A172 cells expressing RFP-tagged mitochondria for 24 to 48 hours. Co-culture was performed either in direct contact or indirectly using 3 μm inserts. Mitochondrial transfer was assessed qualitatively by fluorescence microscopy and quantitatively by flow cytometry (**Figure 1A**).

**Figure 1:**
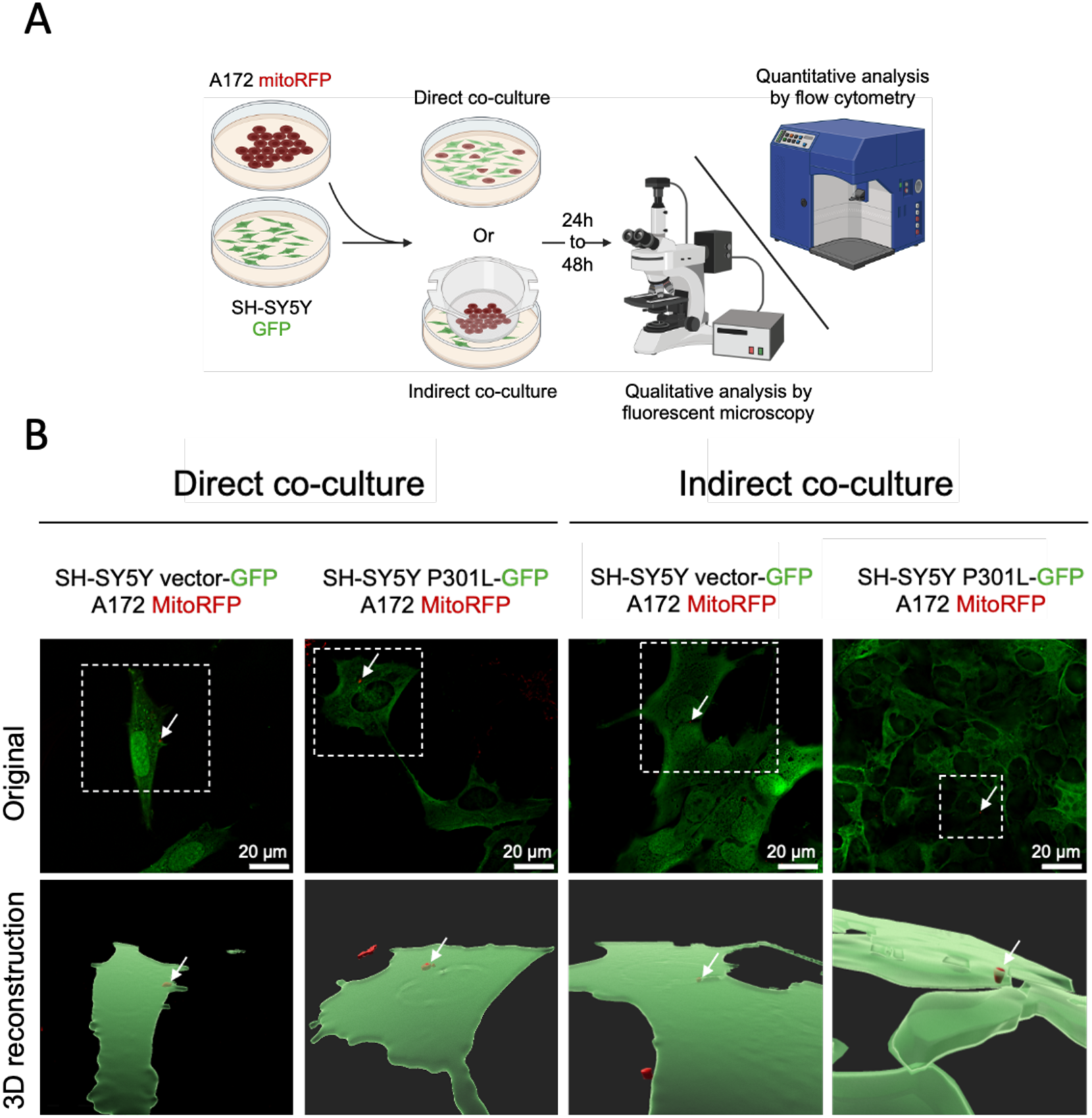
Qualitative assessment of mitochondrial transfer. **(A)** Experimental overview: To investigate mitochondrial transfer from astrocytic A172 cells expressing RFP-tagged mitochondria (A172 mitoRFP) to GFP neuronal SH-SY5Y cells (SH-SY5Y GFP), co-culture experiments were performed either directly (1:1 ratio in the same dish) or indirectly using a 3 μm mesh insert. Mitochondrial transfer was assessed qualitatively by fluorescence microscopy and quantitatively by flow cytometry. **(B)** After 24 hours of co-culture, mitoRFP from A172 cells were detected in GFP-positive SH-SY5Y cells expressing the P301L mutation as well as in vector control cells, in both direct and indirect co-culture conditions (original pictures, top panels). The presence of transferred mitochondria within recipient cells was further confirmed through 3D reconstruction using Imaris software (3D reconstruction, bottom panels).

After 24 hours of co-culture, we visualized a mitochondrial transfer from astrocytes to neurons in both healthy and pathological conditions. The transfer occurred in both direct and indirect co-culture settings, suggesting that at least some of the transfer might occur through extracellular vesicles. A 3D reconstruction using the Imaris software was performed to confirm that RFP-tagged mitochondria were indeed internalized within SH-SY5Y cells rather than being surface-bound (**Figure 1B**).

### P301Ltau increases the mitochondrial transfer from astrocytes to neurons

Since IMT occurs in both healthy and pathological conditions and seems to involve multiple transfer pathways, we aimed to quantify the proportion of mitoRFP in our SH-SY5Y GFP-positive cells using either a contact-dependent or a non-contact-dependent pathway and assess the impact of abnormal tau protein on the transfer (**Figure 2A**).

**Figure 2:**
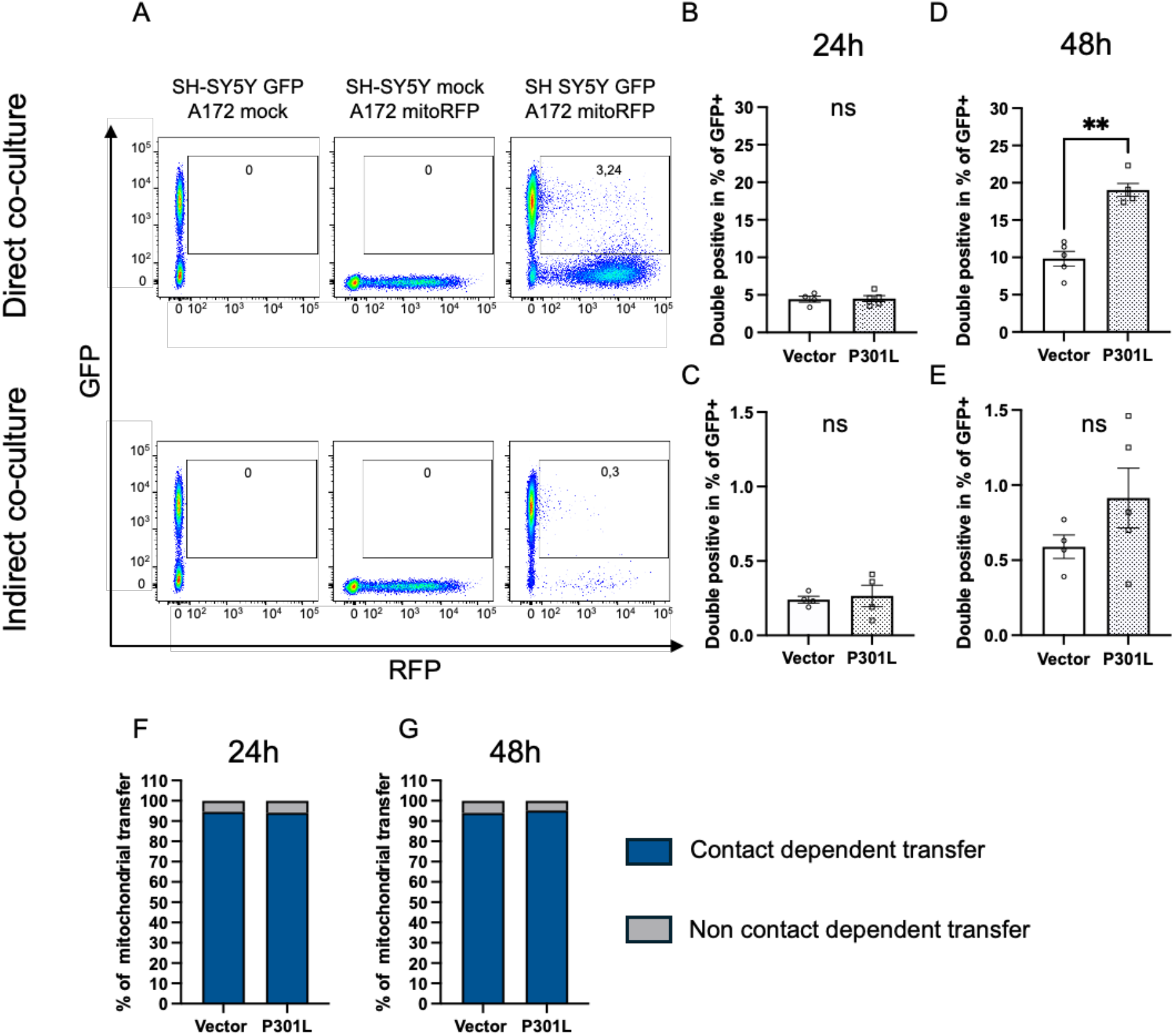
Quantitative assessment of mitochondrial transfer. **(A)** Representative dot plots illustrating the gating strategy for both direct (top) and indirect (bottom) co-culture conditions. MitoRFP-positive events within the GFP-positive SH-SY5Y population (top right quadrant) indicate mitochondrial transfer from A172 astrocytes. **(B-C-D-E)** Bar graphs showing the percentage of GFP-positive SH-SY5Y cells that received astrocytic mitoRFP after 24 and 48 hours of direct or indirect co-culture. **(F-G)** Bar graphs comparing the proportion of mitochondrial transfer occurring via contact dependent versus non-contact-dependent pathways after 24 and 48 hours of co-culture. Data were presented as mean ± SEM. Statistical significance was assessed with Mann-Whitney test. *p* < 0.01 (**), n=4-5. ns= non significant.

IMT quantification by flow cytometry revealed that after 24 hours of direct co-culture, the proportion of mitochondrial transfer from A172 to SH-SY5Y was comparable between vector and P301L-expressing cells, with 4,4% and 4,5% of neurons containing astrocytic mitochondria, respectively (**Figure 2B**). In the indirect transfer setting, only 0,24 % of SH-SY5Y vector cells and 0,26% SH-SY5Y P301L cells contained astrocytic mitochondria, with no statistical difference between the vector and the P301L cells (**Figure 2C**). These results highlight that the contact-dependent transfer appeared to be the predominant pathway, as indirect transfer accounted for only 5,4% (for SH-SY5Y vector cells) and 5,9% (for SH-SY5Y P301L cells) of the total IMT from astrocytic cells to neuronal cells (**Figure 2F**).

After 48 hours of direct co-culture, IMT from A172 to SH-SY5Y was significantly increased under pathological conditions, with 19,0% of neurons containing astrocytic mitochondria in the SH-SY5Y P301L, compared to 9,8% in the vector group (**Figure 2D**). A similar trend was observed in the indirect transfer condition, with 0,91% of IMT detected in P301L-expressing SH-SY5Y cells, compared to 0.59% in vector control cells (**Figure 2E**). Again, the contact-dependent pathway remained predominant in both cell lines after 48 hours co-culture, accounting for 94% of IMT in SH-SY5Y vector cells and 95,8% in SH-SY5Y P301L cells (**Figure 2G**).

### Fates of the transferred mitochondria in recipient cells

Since IMT from astrocytic to neuronal cells increases in the presence of abnormal tau protein, we aimed to determine the fate of the transferred mitochondria in recipient cells. Based on quantification results, we conducted a 48-hour direct co-culture of SH-SY5Y cells expressing both GFP (with or without the P301L mutation) and mitoRFP with A172 cells expressing mitoBFP2. Lysosomes were stained to assess whether donor mitochondria were degraded in recipient cells rather than integrating into the mitochondrial network.

Colocalization analysis revealed that approximately a quarter of the transferred mitochondria fused with the SH-SY5Y mitochondrial network and did not appear to be affected by the presence of abnormal tau protein (**Figure 3B, C**). However, we observed a trend toward higher mitochondrial degradation in the healthy control condition, with 33,4% of transferred mitoBFP2 colocalizing with LysoTracker, compared to 26,7% in the pathological condition (**Figure 3A, C**).

**Figure 3:**
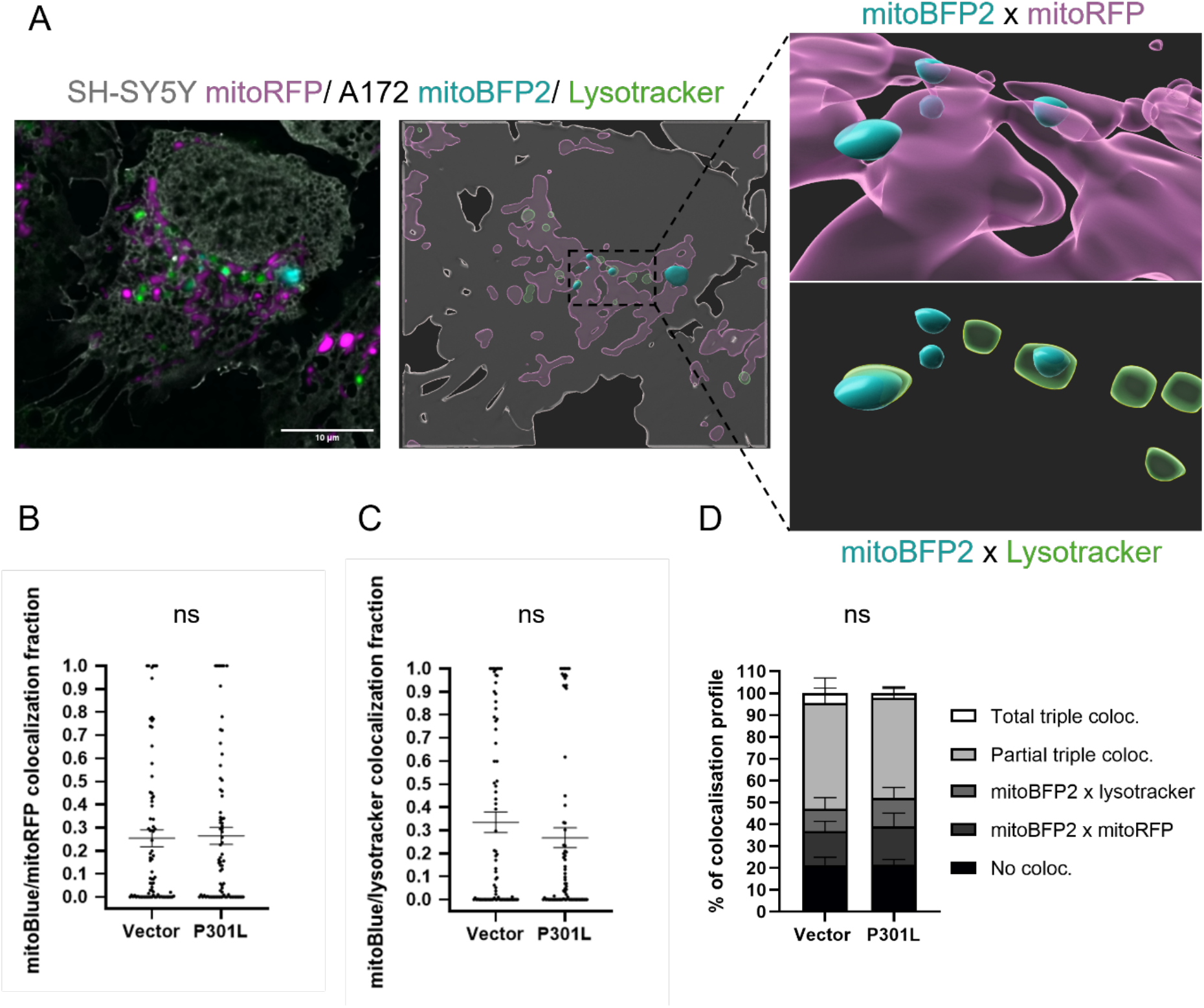
Assessment of donor mitochondrial fate in recipient cells. **(A)** Representative image and 3D reconstruction illustrating the fate of transferred mitochondria (false-colored). SH-SY5Y cells are shown in gray, with their endogenous mitochondria in magenta. Donor mitochondria from A172 cells are shown in cyan, and lysosomes stained with LysoTracker appear in green. **B)** Scatter plot showing the fraction of donor mitochondria (A172 mitoBFP2) colocalizing with the SH-SY5Y mitochondrial network (mitoRFP). **(C)** Scatter plot showing the fraction of donor mitochondria (A172-mitoBFP2) colocalizing with lysosomes (LysoTracker) in SH-SY5Y cells. **(D)** Stacked bar charts representing the distribution of donor mitochondria across five distinct fate profiles in recipient cells, based on colocalization with mitochondrial and lysosomal markers. **“No coloc.”** represents the number of SH-SY5Y cells in which mitoBFP2 does not colocalize with either mitoRFP or LysoTracker. **“mitoBFP2 × mitoRFP”** represents the number of SH-SY5Y cells where mitoBFP2 colocalizes exclusively with mitoRFP. **“mitoBFP2 × LysoTracker”** represents the number of SH-SY5Y cells where mitoBFP2 colocalizes exclusively with LysoTracker. **“Partial triple coloc.”** refers to SH-SY5Y cells in which mitoBFP2 partially colocalizes with both mitoRFP and LysoTracker. **“Total triple coloc.”** refers to SH-SY5Y cells in which mitoBFP2 fully colocalizes with both mitoRFP and LysoTracker. Statistical significance was assessed with Mann-Whitney test for B & C graph and two-way ANOVA for graph D . n=3 with a total of 79 replicates for SH-SY5Y vector cells and 77 for SH-SY5Y P301L cells. ns= non significant

Interestingly, we could identify five distinct populations of SH-SY5Y recipient cells based on the colocalization profile of mitoBFP2 from A172 with endogenous mitochondria (mitoRFP) and lysosomes (LysoTracker). In both pathological and control conditions, a similar proportion of SH-SY5Y cells (∼21%) contained mitoBFP2-positive mitochondria that did not colocalize with either marker, suggesting incomplete integration or a transitional state. A subset of cells showed exclusive colocalization of donor mitochondria with mitoRFP (15.6% in vector cells and 17.6% in P301L-expressing cells) or with LysoTracker (10,1% in vector cells and 13,1% in P301L SH-SY5Y cells). Most cells displayed partial colocalization with both markers, indicating a mixed fate of donor mitochondria. This profile was observed in 48.4% of vector cells and 45.7% of P301L cells. A small population of SH-SY5Y cells exhibited complete fusion of donor mitochondria with its own mitochondrial network, followed by full degradation via the lysosome. This fully colocalized profile was more frequent in vector cells (4.6%) than in cells expressing P301L (2,1%), as shown in **Figure 3D**.

Overall, these data indicate that in both vector and P301L cells, the transferer astrocytic mitochondria are able to fuse with the endogenous mitochondrial network and/or are degraded in lysosomes.

## Discussion

Abnormal tau proteins are known to impair both actin filament dynamics and mitochondrial function, including transport [1,4]. To study the effects of abnormal tau protein on IMT, we used SH-SY5Y cells bearing the P301L tau mutation, a cellular model previously characterized in our laboratory, which displays mitochondrial dysfunctions consistent with those described in the literature [1,2,4,30,31]. Based on our previous findings, we hypothesized that abnormal tau also impacts the mitochondrial transfer between cells. Our results revealed that mitochondrial transfer was increased in the presence of abnormal tau, which aligns with literature showing that pathological conditions and cellular stress stimulate intercellular mitochondrial transfer [32,33]. In our model, the non-contact dependent transfer accounted for only a minor fraction of total IMT, suggesting that contact-dependent transfer pathways, such as tunneling nanotubes and gap junctions, may play a dominant role in IMT from astrocyte to neuron.

Some factors could explain the preferential use of TNTs/gap junctions over vesicle-mediated transfer under pathological conditions. Previous studies have shown that in response to cellular stress, injured or energy-deficient cells can emit signals that induce the formation of TNTs, which are initiated by the stressed cell itself toward a neighboring supportive cell [24,34]. This mechanism could provide a more specific, direct, and faster transfer between two cells than with vesicles while also avoiding exposure of the transferred mitochondria to oxidative stress or lysosomal enzymes present in the extracellular environment. Indeed, it has been shown that non-contact-dependent transfer doesn’t only involve vesicular transport but also includes passive release of mitochondrial material into the extracellular space [16]. The ability of stressed or damaged cells to initiate TNT formation as a “help signal” could explain the unexpected increase in IMT observed in our tauopathy model. One can hypothesize that abnormal tau protein indirectly promotes mitochondrial transfer by impairing mitochondrial function, thereby inducing cellular stress and triggering TNTs formation. Future studies will aim to elucidate the underlying mechanisms involved in this process. Given the observed increase in IMT from A172 to SH-SY5Y in the context of tauopathy, we initially hypothesized that a greater proportion of transferred mitochondria would integrate into the mitochondrial network of SH-SY5Y cells expressing the P301L mutation as a compensatory response to tau-induced mitochondrial dysfunction. We managed to classify recipient cells into five distinct populations based on the colocalization results of donor mitochondria with endogenous mitochondria and lysosomes. Surprisingly, the distribution of these profiles was largely similar between pathological and control conditions, suggesting that the presence of abnormal Tau does not entirely alter the fate of transferred mitochondria at the population level. However, we still identified a difference in one specific profile: the proportion of cells that fully integrated donor mitochondria into their mitochondrial network followed by complete lysosomal degradation trended to be higher in vector control cells than in P301L-expressing cells. This may explain why we observed a higher proportion of donor mitochondrial colocalizing with lysosomes in the SH-SY5Y vector condition. A possible interpretation is that mitophagy is impaired under pathological conditions. Since the proportion of cells showing only donor mitochondrial degradation was similar between groups, it suggests that the intensity of degradation may be negatively impacted in the presence of abnormal tau. This aligns with previous studies showing that tauopathies can interfere with mitophagy [15,35]. Moreover, studies have shown that healthy mitochondria transplantation increases mitophagy in recipient cells [36], which can also explain these results. Indeed, since mitophagy is not impacted in vector cells, the transfer of new mitochondria from A172 might lead to a mitophagy-boosting effect. Additional studies will be conducted to confirm or refute this hypothesis.

## Conclusion

Our findings demonstrate that abnormal tau expression enhances mitochondrial transfer from astrocytic to neuronal cells, likely as a response to tau-induced mitochondrial stress. This transfer appears to occur predominantly via contact dependent mechanisms in healthy and pathological conditions. Although the overall fate of transferred mitochondria was similar between pathological and control conditions, subtle differences point toward impaired mitophagy of the donor mitochondria in the recipient cells in the presence of abnormal tau protein. The lack of mechanistic experiments limits our ability to fully understand the causes and pathways underlying the observed increase in IMT; however, addressing this will be a key priority in our future work. Nevertheless, to our knowledge, we are the first to show the impact of tau on IMT between astrocytic and neuronal cells in the context of tauopathies.

## Acknowledgements

This work was supported by grants from the Novartis Forschungsstiftung FreeNovation, the OPO Foundation, the Dementia Research, Synapsis Foundation Switzerland (2022-PI05) and the Bürgenstock Foundation. We thank Fides Meier and the flow cytometry core facility of the department of biomedicine of Basel (Switzerland) for the technical support.

## Author contributions

**Aurélien Riou**: Writing-original draft, Conceptualization, Methodology, Data collection, Formal analysis; **Aline Broeglin**: Methodology, Data collection, Formal analysis, Writing-review; **Amandine Grimm**: Funding acquisition, Conceptualization, Data collection, Methodology, Formal analysis, Writing-review and editing, Resources, Supervision; **Anne Eckert**: Writing-review, Supervision, Resources; **Andreas Papassotiropoulos**: Writing-review, Supervision, Resources All authors have read and agreed to the published version of the manuscript.

